# New Functions Identified for HIV gp120 with the New ProtSub Sequence Matching

**DOI:** 10.1101/2022.02.01.478707

**Authors:** Benjamin Litterer, Sayane Shome, Kejue Jia, Robert L. Jernigan

**Affiliations:** Bioinformatics and Computational Biology Program, Biophysics and Molecular Biology, Iowa State University, Ames, IA, 50011, USA; Roy J. Carver Department of Biochemistry, Biophysics and Molecular Biology, Iowa State University, Ames, IA, 50011, USA

## Abstract

The reduced cost of sequencing is leading to an explosive growth in the number of available sequences across diverse genomes, and for individual patients. Inferring meaningful functions of individual genes/proteins is lagging, which hinders the deeper understanding of biological function and evolution. Traditionally, protein function has been determined by time consuming experimental methods or by sequence matching that often does not agree with the experimental findings. We have significantly improved protein sequence matching, by accounting for inter-dependent amino acid substitutions observed within densely packed protein structures, which yields additional substitutions beyond those usually seen, with good matches to additional proteins, some having new functions, not identified by conventional sequence matching. In the current study, we have applied this approach to predict novel functions for the proteins from HIV. These newly found functional annotations are then manually reviewed and many are validated from the literature, here for the HIV envelope protein gp120. These new functions are both more specific as well as some being entirely novel functions. We also show statistically that on average our new functional annotations are more informative than those given by conventional substitution matrices such as BLOSUM62. These results suggest that the new ProtSub protein sequence matching that incorporates structural information generally yields better identifications of related proteins, which can have broader and often gains in identifying more specific functions

## Introduction

The Human genome sequence has been followed rapidly by the sequencing of many other genomes, and there are presently sequence available from more than 292,000 proteomes (UniProt, 2021). In addition, the cost of obtaining genome sequences has plummeted to the point where sequencing individual patients is feasible, so already there are more several hundred million protein sequences. The protein sequence database, UniProt Knowledge Base, currently has over 189 million proteins, a major increase from the 120 million reported in the UniProt Consortium’s 2019 publication (UniProt, 2021; UniProt, 2019). Despite these gains in the number of sequences, some fraction of newly sequenced proteins do not have annotated functions, and due to the expense of experimentally obtaining functional information, it enable doing this computationally. Also it should be noted that there is an open competition, CAFA (Zhou et al., 2019) with the specific aim of improving and enhancing function predictions.

The most common approach to making assignments of function is by sequence matching, which can be seen among high-performing functional predictions (Zhou et al., 2019). Traditionally, proteins with sufficiently similar sequences are assumed to have similar structures and similar functions. This enables the inference of function between proteins having sufficiently similar sequences. This often works well for many cases, with the exceptions being few cases where the same function is found in extremely dissimilar structures, which have sequences lying in the “twilight zone” with respect to one another.

Sequence matching most commonly relies on the use of amino acid similarity matrices, a 20×20 symmetric table that shows how similar two amino acids are to one another. And for example, BLAST as installed for public use at NCBI and the National Library of Medicine utilizes the series of BLOSUM Matrices derived by the Henikoffs (Henikoff & Henikoff, 1992). But, there are aspects of structure that do not let similar amino acids to always be easily substituted for one another. The dense packing inside of globular proteins means that multiple substitutions can occur that do not strictly conform to substitutions of similar amino acids in each individual position in a structure. There can be compensation. The simplest example of such dependences are pairs where the individual amino acids are substituted for one another. These are common when the pair is directly interacting; these can be substitutions such as a small-large pair being changed to a large-small pair or a +/− pair being changed to a −/+ pair.

To include such structural information in sequence alignments, we have previously developed a new substitution matrix, ProtSub, which accounts for such interdependences (Jia & Jernigan, 2021), so it is not simply an amino acid similarity matrix but goes beyond that by utilizing the concept that the pair substitutions within a pair of coevolved residue sites contain important information about the structure and function. These coevolved sites can be identified through computational approaches and the interdependent substitutions within those sites can be employed to derive a matrix describing the likelihood of changes between pairs of specific amino acid types. Since ProtSub does not penalize sequence changes that preserve structure, it allows more amino acid substitutions. This brought into agreement cases that had high levels of structure similarity but previously did not have sequences that were similar enough to indicate the structural similarity. ProtSub exhibits significantly better agreement between the sequence alignment and the structure alignment of proteins so that residues aligned in the sequence match are close together in the structure match, in comparison with other substitution matrices (Jia & Jernigan, 2021). Here we exploit this in a practical application for the functional annotations of the HIV proteins, which is particularly appropriate because it is a rapidly evolving virus and thus has large number of available sequences, permitting the generation of reliable multiple sequence alignments.

HIV is a viral infection that weakens the immune system through gradual death of CD4+ T cells. Infection is associated with cardiovascular and bone disease as well as a greater risk of infections and cancers throughout the body. HIV primarily infects lymphocytes, monocytes, macrophages, and dendritic cells (Deeks, Overbaugh, Phillips, & Buchbinder, 2015). Cell entry is initiated when the gp120 protein of the HIV envelope (env) protein binds to the CD4 receptor of the target cell. This triggers a conformational change in gp120, allowing it to bind to either the CCR5 or the CXCR4 coreceptors, depending on the strain. The second component of the HIV envelope, gp41, then initiates membrane fusion, resulting in eventual entry of the virus into the host cell (Kovalevich & Langford, 2012). Given its crucial role in HIV pathogenesis, this protein and its effects are widely studied, and it was of great interest in analyzing the output of our method. Although antiretroviral therapies (ART) are extremely effective in prolonging life, HIV remains a pertinent global public health issue with nearly 1 million deaths per year and multiple countries having prevalence rates above ten percent in 2017 (James et al., 2018).

## Results

When comparing functional assignments made with the BLOSUM62 and ProtSub matrices, we find that ProtSub identifies significantly larger numbers of functions and that these include most of those in with the BLOSUM62 output (Table 1). Across all HIV proteins, we find that ProtSub assigns 1,831 more functions than BLOSUM62 and overlaps with 75% of the functions assigned using BLOSUM62 (see Table 1 for details). This indicates that despite the ProtSub matrix being more permissive in the substitutions that it allows, it still aligns with sequences that may not require this higher order consideration of amino acid interdependencies.

**Table 1.** Summary of the annotations assigned unique with each matrix, as well as annotations which were found with both.

|  |  |  |  |
| --- | --- | --- | --- |
| HIV Protein | Unique to ProtSub | Unique to BLOSUM62 | Both Matrices |
| gagPol | 389 | 16 | 28 |
| vpu | 44 | 0 | 8 |
| vpr | 64 | 0 | 14 |
| virion | 148 | 1 | 5 |
| nef | 157 | 10 | 15 |
| env | 498 | 4 | 19 |
| rev | 112 | 0 | 6 |
| gag | 405 | 14 | 25 |
| tat | 60 | 1 | 18 |
| **Total:** | **1877** | **46** | **138** |

Our review of the assigned GO terms shows that many of the functions found with the ProtSub matrix are more specific and more informative than those from BLOSUM62. To demonstrate this, the information content was computed for each function identified. The mean information content for the ProtSub matrix was found to be .948 nats^1^ higher than the mean information content of the BLOSUM62 output, and the difference in median information content was 1.066. To determine the significance of this difference, both parametric and bootstrap hypothesis tests were applied. Welch’s test for samples of unequal variance confirms the significance of this value with a highly significant p-value below 0.001. For the bootstrap test, the null distribution was created by drawing with replacement from the combined information content values for annotations from both matrices. The difference in sample means and medians was then calculated, and the process repeated 5000 times. The observed differences in sample means and medians was sufficiently large that it was not found in the null distribution created with the bootstrapping procedure. These results indicate that annotations found with the ProtSub matrix carry significantly more information than those found with BLOSUM62, which is quite obvious for many of the cases.

While the distribution of information content measures the overall gain in specificity of our new functions, it does not demonstrate the direct gains for any particular function with reference to what was found using BLOSUM62. To show this, we use the GO hierarchy to find the closest function assigned with BLOSUM62 for each of our newly assigned functions found with ProtSub. From this comparison, we find that 88.4% of our new annotations are more specific than their nearest neighbor in the BLOSUM62 set (Fig. 1).

**Figure 1.**
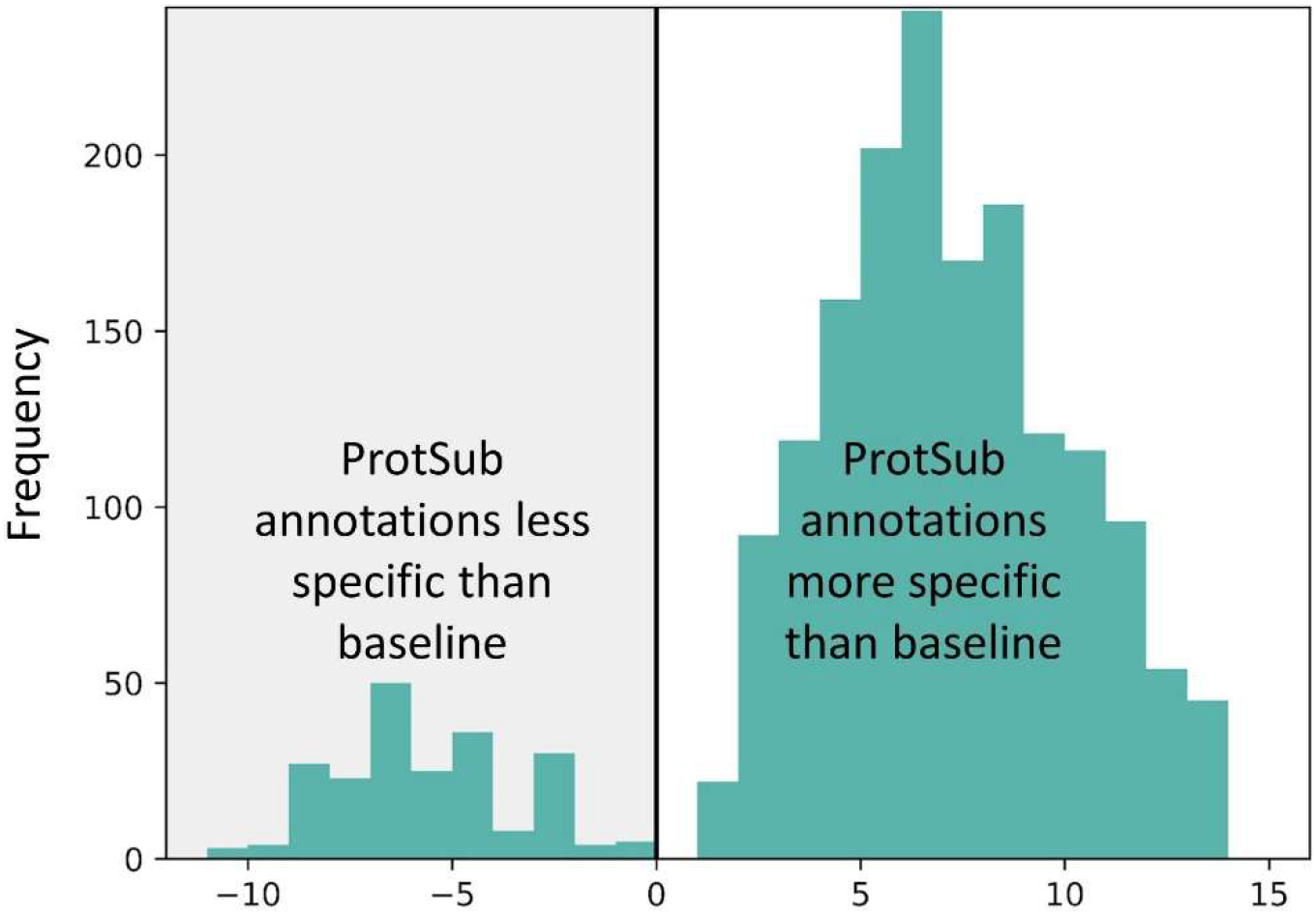
The distance between each new function assigned using the ProtSub matrix to its closest annotation neighbor assigned using the BLOSUM62 matrix (termed baseline here). On the left are shown the cases that are less specific and on the right ones that are more specific. Distance is defined as the sum of the distances from each annotation to their deepest common ancestor. Clearly the gains outweigh the losses.

In the comparison made, it is not clear whether the comparisons are being made between related functions or not. To aid in the further interpretation of our results we next cluster the functions found into related groups by choosing those similar to one another. These discrete functional clusters can then be assessed among the functions within the cluster. Doing this reduces the 2,015 output protein functions from the new coevolution-based approach to 451 clusters spanning the three GO sub-ontologies and separately for the nine different proteins. We then keep only the clusters that include a function assigned with BLOSUM62 and compared the median information content for the ProtSub and BLOSUM62 annotations. As shown in Fig. 2, annotations from our matrix were equally or more informative for 84.3% of the clusters, a result that is not very different from the first comparison made above

**Figure 2.**
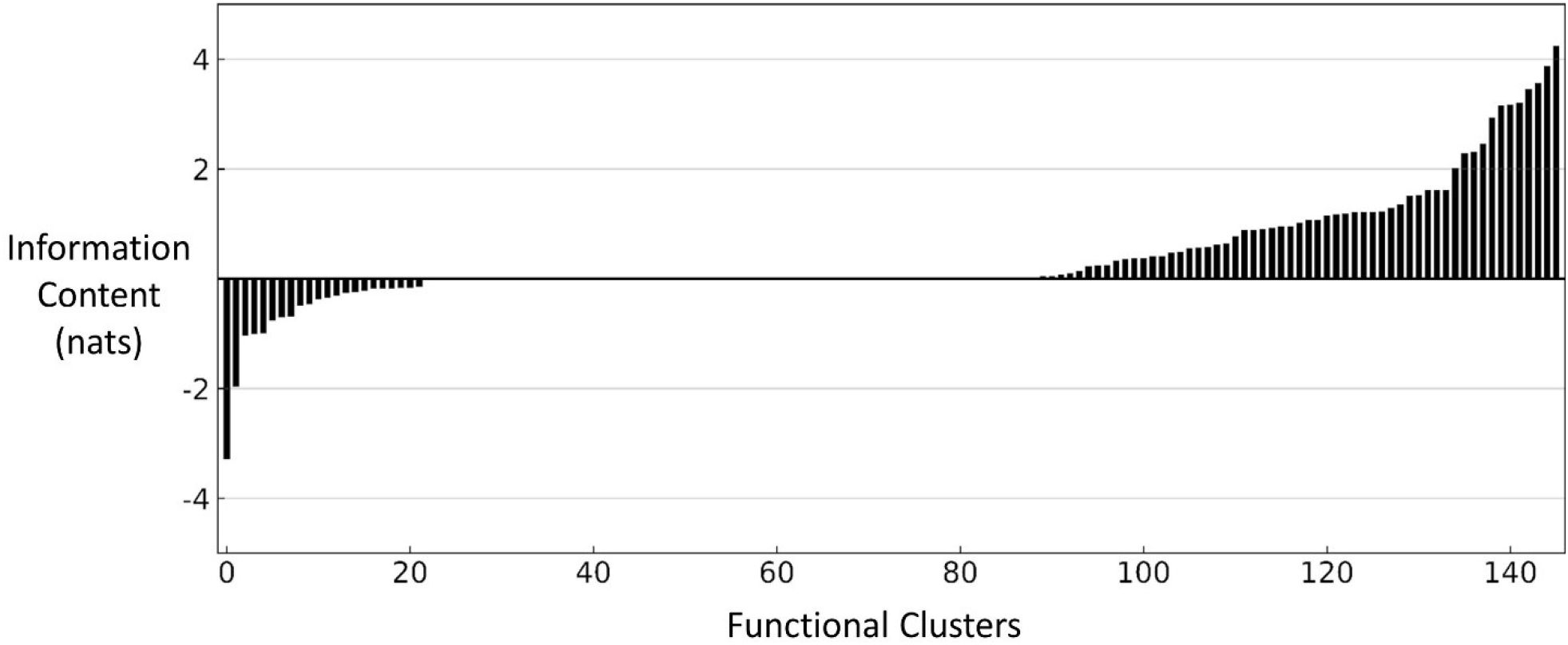
Information Content Differences within Functional Clusters: The difference in median information content between output with ProtSub and the output with BLOSUM62 for each functional cluster.

The dendrograms in Fig. 3 show that the output can be classified primarily into two groups. Many of the functions were similar to the output given from using the BLOSUM62 matrix but were much more specific or added related information. As shown in Figure 3.B, the “actin filament reorganization” function was given by the output when using both the BLOSUM62 and the ProtSub matrix; however, output from the ProtSub matrix gave significantly more annotations all relating to cytoskeletal activity and the broader processes that are facilitated by it such as synapse formation. Another example of this type of information gain is shown in Figure 3.A, which includes many additional annotations indicating the particular mechanisms related to the function “evasion or tolerance by virus of host immune response.” In addition, much of the output unique to the ProtSub matrix results gives insight into areas of protein function that were not indicated in the BLOSUM62 results. One such case is shown in Figure 3.C, where the new functions from ProtSub describe the potential role of the HIV envelope in development. No output relating to this function was given by output from the BLOSUM62 matrix.

**Figure 3.**
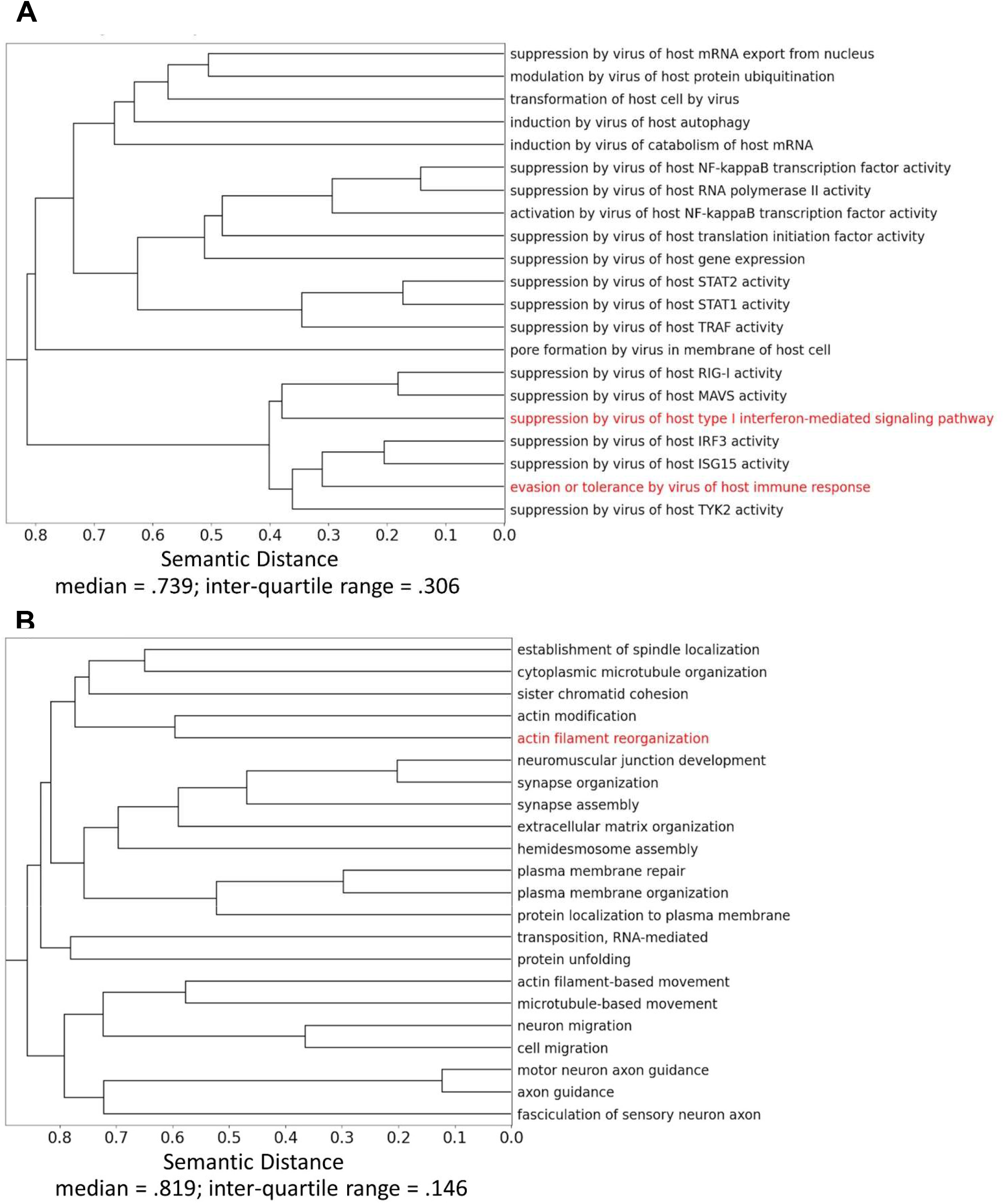
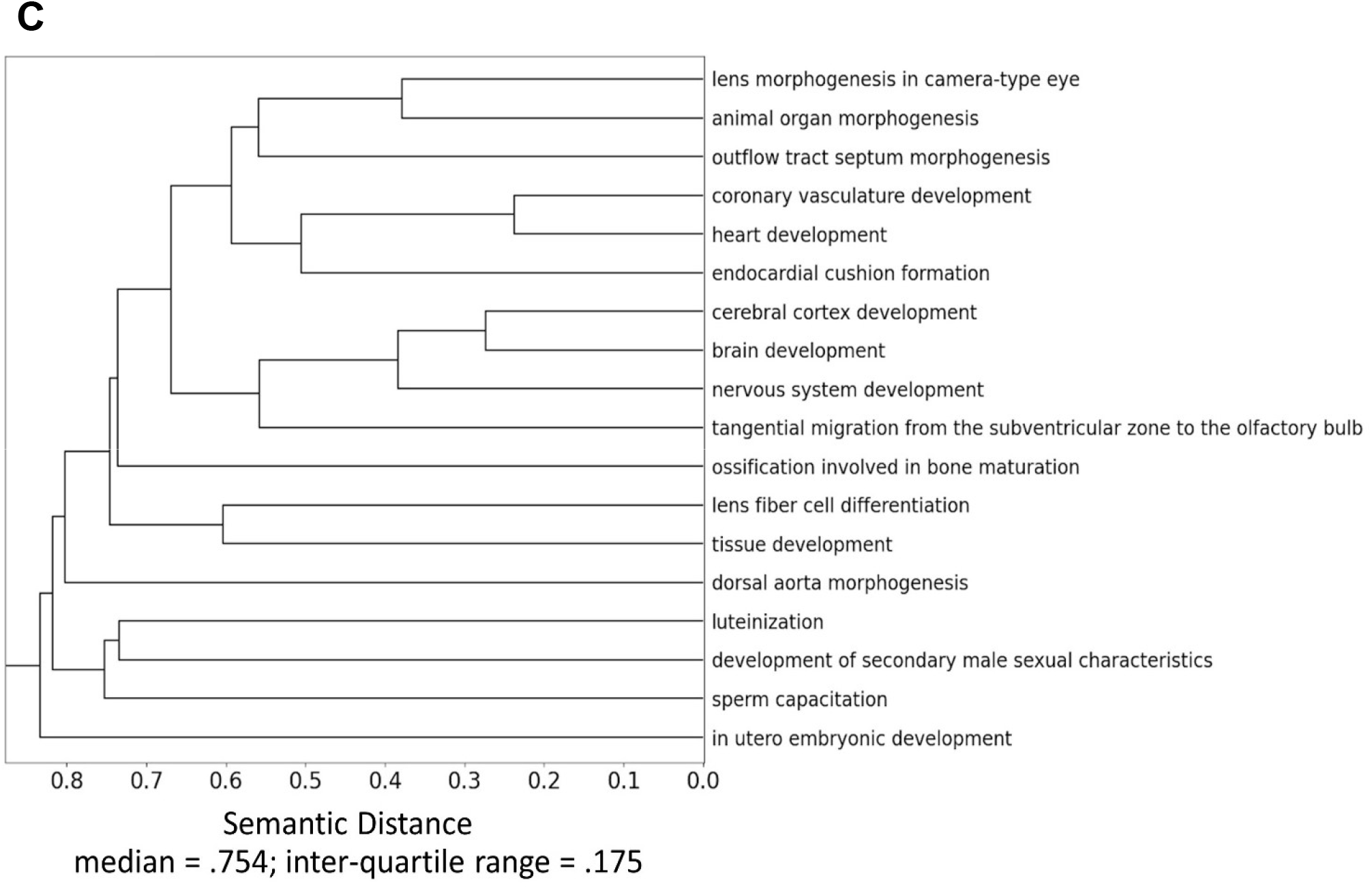
Dendrograms show the results of the calculated semantic distances (1 – GOGO similarity) between GO terms and using hierarchical clustering with average linkage. Part A,.B, and C all depict various subtrees of a larger dendrogram for the Biological Process annotations of the HIV envelope given by ProtSub. The three functions in red are those also obtained with the BLOSUM62 matrix. Annotations found by BLOSUM62 but not ProtSub were “‘suppression by virus of host tetherin activity”, and “membrane fusion involved in viral entry into host cell”. Median and inter-quartile range of the entire tree (some annotations not depicted in 3.A, 3.B, 3.C) are 0.982 and 0.034 respectively. These statistics were found from the distribution of the distances between every pair of annotations in the given tree or subtree.

## Discussion

### Validation of New Functions

In this section we compare some of the functions that have been identified here with literature reports of functions. This is our validation procedure, and the results are divided into several categories of functions, indicated by the headers of the following sections. A summary of these experimentally-validated functions is found in Table 1. Additionally, functions related to the groupings below but still needing experimental validation are shown in Supplemental Tables 1 and 2, that are for p-values of less than .001 and greater than .001, respectively.

**Table 2.** New Functional Annotations and Supporting Evidence for 4 Broad Classes.

|  |  |  |  |
| --- | --- | --- | --- |
| Function |  | Citation(s) | p-value (Fisher's test) |
| **1. Neuron/Axon Effects** | 1. learning or memory | (Jaeger & Nath, 2012) | .0008 |
|  | 2. startle response | (Fitting et al., 2006) | .0052 |
|  | 3. microtubule-based movement | (Bachis et al., 2006; Berth et al., 2015) | <.0001 |
|  | 4. transmembrane receptor protein tyrosine kinase signaling pathway | (Lee et al., 2003) | <.0001 |
|  | 5. chloride transmembrane transport | (Lee et al., 2003) | <.0001 |
|  | 6. cation transport | (Lee et al., 2003) | .0011 |
|  | 7. calcium ion import | (Benos et al., 1994) | .0365 |
|  | 8. intracellular pH elevation | (Benos et al., 1994) | <.0001 |
|  | 9. calcium ion transmembrane transport | (Benos et al., 1994) | <.0001 |
|  | 10. potassium ion transmembrane transport | (Benos et al., 1994; Chen et al., 2011; Lee et al., 2003) | .0011 |
|  | 11. release of sequestered calcium ion into cytosol | (Benos et al., 1994) | <.0001 |
| **2. Glioblastomas, Localization, Migration and Adhesion** | 1. regulation of cell migration | (Valentin-Guillama et al., 2018) | <.0001 |
|  | 2. isoleucyl-tRNA aminoacylation | (Valentin-Guillama et al., 2018) | <.0001 |
|  | 3. leucyl-tRNA aminoacylation | (Valentin-Guillama et al., 2018) | <.0001 |
|  | 4. valyl-tRNA aminoacylation | (Valentin-Guillama et al., 2018) | 0.0003 |
| **3. Synapse and Fusion** | 1. synapse assembly | (Jolly, 2010) | .0001 |
|  | 2. synapse organization | (Jolly, 2010) | .0001 |
|  | 3. positive regulation of synapse assembly | (Jolly, 2010) | <.0001 |
|  | 4. protein localization to plasma membrane | (Jolly, 2010) | .0138 |
|  | 5. cell-cell adhesion | (Ballana et al., 2009) | <.0001 |
|  | 6. cell adhesion mediated by integrin | (Ballana et al., 2009) | .0001 |
|  | 7. cell-matrix adhesion | (Ballana et al., 2009) | .0358 |
| **4. Development** | 1. brain development | (Nozyce et al., 1994) | <.0001 |
|  | 2. cerebral cortex development | (Nozyce et al., 1994) | .0215 |
|  | 3. nervous system development | (Nozyce et al., 1994) | <.0001 |
|  | 4. in utero embryonic development | (Dibbern et al., 1997) | <.0001 |
|  | 5. tissue development | (Dibbern et al., 1997) | <.0001 |

#### Neuron/Axon Activities

There have been numerous reports of the effects of the HIV on the brain and central nervous system. These include conditions such as HIV associated neurocognitive disorder and HIV associated dementia with manifestations in recall and memory, difficulty with verbal tasks, reduced problem solving ability and a host of physical effects such as slower movements of the eyes and limbs, and an unsteady gait (Price et al., 1988). Other cognitive effects include alteration in the perception of sound, as evidenced by studies in both mice and humans reporting changes in startle response following an initial auditory stimulus (Fitting, Booze, & Mactutus, 2006; Minassian et al., 2013). Ultimately, HIV’s infection of the Central Nervous System (CNS) may progress to severe dementia, with patients being in an almost “vegetative state”, unable to speak or walk. Remarkably, the new ProtSub matrix yields statistically significant functional annotations corresponding to several of these clinical observations. Our results include annotations not found by BLOSUM62 such as “learning or memory”, “neuromuscular process controlling balance”, and “startle response” (see Table 2, items 1.1, 1.2; Supp. Table 1, item 1.1).

While the correspondences between these symptoms reported from clinical studies and the functions identified with our computational method are striking, these annotations are all assigned specifically to the HIV Envelope (gp160) protein, which is remarkable that one protein could cause so many profound effects. However, gp160 is further cleaved into gp120 and gp41, and there is a wealth of experimental evidence confirming these linkages with gp120. Since HIV is known to pass the blood brain barrier (McRae, 2016), infected macrophages and monocytes are believed to be the source of gp120 within the central nervous system. The entry of these cells into the CNS is facilitated by a gp120-dependent increase in the expression of the intracellular adhesion protein ICAM-1 (Shrikant, Benos, Tang, & Benveniste, 1996). These infected cells shed gp120, which has been found to cause neuronal apoptosis and degeneration in transgenic mice models (Jaeger & Nath, 2012). In addition, damage to the blood brain barrier allowing HIV to freely enter the central nervous system has also been shown to be affected by gp120. This damage to neurons from gp120 underlies the neurological effects seen in HIV associated dementia (Kanmogne, Kennedy, & Grammas, 2002). Once gp120 breaches neurons, its spread is facilitated by transport along motor-dependent microtubules (Bachis, Aden, Nosheny, Andrews, & Mocchetti, 2006; Berth, Caicedo, Sarma, Morfini, & Brady, 2015). Both the presence of gp120 within the central nervous system, and its transport along axons via microtubules are confirmed by the functional annotations from ProtSub.

There are multiple experimentally observed mechanisms by which HIV causes cell death once it is within the CNS. Changes to microtubules after gp120 binding have been reported to result in “acute or chronic axonal and dendritic fragmentation” (Avdoshina, Taraballi, Tasciotti, Uren, & Mocchetti, 2019). Another main area of previous research focus has been on the regulation of ion flow in neurons, mitochondria, astrocytes, and macrophages. In one study, gp120 was reported to alter the outward potassium currents leading to neural cell death in “rat cortical neuronal cultures” (Chen et al., 2011). In another study, HIV was reported to alter the membrane potential of mitochondria, which could contribute to the dysfunction of neural cells (Teodorof-Diedrich & Spector, 2018). In astrocytes, gp120 initiated a change in the Na+/H+ exchange “mediated by activation of a tyrosine kinase”. This resulted in rising levels of potassium ions around neural cells and an increase in calcium levels inside neurons, which was hypothesized to lead to neuron dysfunction (Benos, McPherson, Hahn, Chaikin, & Benveniste, 1994). Additionally, gp120 was said to be detrimental through its binding to chemokine receptors found on macrophages. This alters the flow of potassium and chloride ions, increasing the levels of cellular calcium ions, and activates a tyrosine kinase, all of which seems likely to contribute to the progression of HAD (Lee et al., 2003). These experimental findings correspond to the new functional assignments (Table 2, items 1.3, 1.4, 1.5, 1.6, 1.7, 1.8, 1.9, 1.10, 1.11).

After reviewing HIV pathogenesis literature, we can confirm that our computationally derived functional annotations accurately describe the widely reported roles of gp120 in HIV neurocognitive disorder and HIV associated dementia. These annotations are not included in the known Gene Ontology annotations, and neither were they produced by the BLOSUM62 results. This is only one set of cases demonstrating the ability of the new amino acid substitution matrix to make sequence matches to a wider range of proteins with more specific functions that clearly closely relate to the HIV disease manifestations. This suggests that judicious application of the new approach can reliably produce functional results of diseases in other cases that were only found by costly and slow experimental methods, and could therefore also help to point out more appropriate directions for investigation significantly more effectively

#### Glioblastomas and cell migration

Another problem associated with HIV is an increased likelihood of the development of Glioblastoma tumors. Gp120 has been reported to contribute to the development of these tumors shown in both *in vivo* and *in vitro* experiments. In particular, gp120 has a positive effect on cell migration and proliferation in experimental assays. Downregulation of amino acids including valine, leucine, and isoleucine has also been reported, in addition to changes in aminoacyl-tRNA biosynthesis (Valentin-Guillama et al., 2018). These experimental results correspond to functions assigned using the ProtSub matrix (Table 2, items 2.1, 2.2, 2.3, 2.4), and there are additional migration-related annotations requiring further study and analysis (Supp. Table 1 item 2.1, Supp. Table 2, item 2.1). Chemokines are thought to be involved in migration in the brain, and it has been hypothesized that viruses such as HIV binding to chemokine receptors could interfere with this role. This could contribute to the previous discussion of neurological dysfunction associated with HIV infection (Tran & Miller, 2005). To our knowledge, more research in this area is necessary to fully confirm these new annotations.

#### Synapse and Fusion

Virological Synapse is a process wherein a molecular structure forms between an infected cell and a non-infected cell for cell fusion. HIV is reported to form such structures between infected T cells and dendritic cells, infected T cells and macrophages, and between infected T cells and uninfected T cells. Normally T cells would not form long term contacts with other T cells, but the stability of this interaction when one is infected with HIV is driven by the Env for T cells binding to other T cells. In particular, gp120 binding is theorized to activate T cell receptor signaling, resulting in polarization of the microtubule organizing center, actin polymerization, and an F-actin depleted zone (Vasiliver-Shamis, Cho, Hioe, & Dustin, 2009). Once the synapse is formed, Env, Gag, and lipid rafts move towards the synapse due to polarization of the microtubule organizing center (Jolly, 2010). Synapse maintenance and promotion is also aided by the presence of intracellular adhesion molecules such as ICAM-1. Gp120 is known to increase the expression of this molecule in endothelial cells, which results in enhanced infectivity (Ren, Yao, & Chen, 2002). The contribution of the HIV envelope protein to virological synapses was first discovered experimentally, but it is also confirmed in our computationally predicted annotations (Table 2, items 3.1, 3.2, 3.3, 3.4). Although they are not linked to virological synapse formation, additional research has confirmed our annotations related to cell-matrix adhesion (Table 2, items 3.5, 3.6, 3.7), as well as similar annotations that require some further confirmation (Supp. Table 1, items 3.1, 3.2, 3.3).

#### Development

Although HIV is transmitted from one adult to another, there is also significant transmission in the uterus and during childbirth. This problem is particularly significant in countries where childbirth commonly occurs outside of health facilities (Mwaba, Ngoma, Kusanthan, & Menon, 2015). Intrauterine and pediatric HIV infection results in a set of developmental problems, in addition to the symptoms common in HIV positive adults. Abnormalities in both cognitive and physical development occur through a variety of mechanisms. The birth weight and size of HIV positive babies is known to be below normal, which is a result of delayed development in the uterus. Gp120 may be a direct link to this observation, as the presence of gp120 in mice embryos at very low concentrations is reported to stunt growth (Dibbern, Glazner, Gozes, Brenneman, & Hill, 1997). In the case of delayed postnatal neurodevelopment, it is likely that the effects of gp120 on the adult brain may also be seen in infants and cause cognitive abnormalities for children (Nozyce et al., 1994). These experimental and clinical findings for HIV effects on development also are confirmed in our results (Table 2, items 4.1, 4.2, 4.3, 4.4, 4.5). Our results also include a set of annotations related to development that do not appear to have been experimentally confirmed and these could indicate new directions for further research (Supp. Table 1 items 4.1, 4.2; Supp. Table 2, items 3.1, 3.2, 3.3).

## Conclusions

Using the ProtSub matrix in protein sequence matching significantly enlarges and improves upon the functional assignments made from conventional sequence matching. This has been demonstrated here where our computational analyses demonstrate the statistical significance and information content of the new annotations. In addition, a set of manually reviewed functions, primarily involving gp120, are found to be confirmed in the experimental HIV literature. Given these results, there is a significant need to extend the application of our method more widely. Doing so should provide a substantial increase in putative protein functions, particularly relating to disease and organismal malfunction. To scale up this approach requires reliable information on the accuracy of our assignments as compared to a ground truth dataset, such as those available from the CAFA challenge (Zhou et al., 2019). Using such a dataset would allow for a statistical analysis of the quality of our annotations and a comparison of our method with other approaches. We are planning to carry out such a study.

## Methods

### Dataset collection

We collected protein sequences for the nine HIV proteins given in the UniProt reference proteome database by choosing an HIV strain having the most complete subtype indicated by UniProt’s Complete Proteome Detector Algorithm. The next step involved finding the Gene Ontology (GO) terms currently associated with these protein sequences. A tabular file was downloaded from the UniProt web site that contains cross references from Swiss-Prot IDs to GO annotation terms. It should be noted that these cross-reference files contain the majority of assigned functions, but some have been removed by SwissProt’s filtering algorithm, which has a preference for manually, rather than electronically assigned protein functions. The SwissProt database was chosen instead of other larger databases because it has been manually curated, providing some higher level of confidence in its functional assignments. The HIV sequence and functional assignments for the SwissProt database were the starting point for the rest of the analysis.

### New potential homologs are found with BLAST using the new ProtSub substitution matrix

The BLAST algorithm (Altschul, Gish, Miller, Myers, & Lipman, 1990) is used to identify the sequence hits for each input query sequence based on a set of parameters specified. For carrying out a comparative performance analysis of ProtSub with commonly used scoring/substitution matrix BLOSUM62, we configured the BLAST software with these two matrices in separate runs. We queried all the nine HIV sequences using the above-mentioned two different builds of BLAST to find the sequence hits in the Swiss-Prot database (Bairoch & Apweiler, 2000) with an e-value cutoff of .001. We then make assignments of function by using all the sequence hits and their associated GO IDs. These IDs are part of a set of controlled vocabulary that represent the different functions, processes, and cellular components that a given protein may take part in. Annotating the sequence matches with their respective GO IDs provides a set of candidate functions (IDs) that could be potentially transferred between the query and other matching sequences.

To determine which candidate functions occur significantly more often than random chance, Fisher’s exact test is performed by comparing the frequency of a given function in the output to the frequency of that function in the annotations of the Swiss-Prot database (Bairoch & Apweiler, 2000). This has the effect of comparing a function’s frequency to the expectations from random sampling from the Swiss-Prot database. Functions having a p-value less than .001 are attributed to the query protein on this basis. This results in each HIV protein having two sets of computationally predicted functions corresponding to the two sequence substitution matrices that were used. Next, we compare the two sets of results. We find a large number of additional functions with the new ProtSub matrix that we have developed (Jia & Jernigan, 2021).

### Comparison of ProtSub results with the BLOSUM62 results

For a given set of functions assigned to a query protein, we found the distance in meaning (semantic similarity) between GO terms using the GOGO package (Rivals, Personnaz, Taing, & Potier, 2007). Since GO terms are arranged in an ontology from least specific to most specific, GOGO leverages the structure of this ontology to determine how similar two functions are. Although there are many ways to define semantic similarity in the Gene Ontology, there are two common caveats. First, an equal distance in semantic meaning is assumed for all edges in the gene ontology. GOGO addresses this concern by using weighted edges. Second, bias is introduced when using methods that simply count the frequency of annotations for a large set of proteins and derive information content from this frequency. GOGO addresses this concern by calculating the informativity of a GO term using the number of children of its ancestors in the GO hierarchy, which is not biased by the amount of study and subsequent annotation of a particular function. Semantic distances calculated with this method were used to carry out hierarchical clustering with results demonstrated in Figure 3. Each query protein had three associated dendrograms, one for each of the three main sub-ontologies in the GO: Molecular Function, Cellular Component, and Biological Process.

To determine how much information a given functional annotation contains for a query protein, we use information content, which is calculated by taking the negative natural log of the probability of randomly selecting the GO term in the Swiss-Prot database. This probability is calculated by using the Swiss-Prot-GO cross reference file noted above in the <u>Dataset Collection</u> section.

The GOGO similarity (Zhao & Wang, 2018) was evaluated to calculate distances between functions, and these functions were then clustered using the graph-based affinity propagation algorithm from the sci-kit learn package. This is a simple way to classify and understand the new functions found. A full list of clustered annotations found with ProtSub are compiled in Supplemental File 1. In particular, we clustered annotations in the ProtSub results, and noted which clusters had annotations also in the BLOSUM62 results. This allowed us to determine the difference in mean and median information content for annotations from the two results within each cluster.

In addition to considering functional clusters, we computed the distance in the GO hierarchy between each new annotation found to its closest annotation found by BLOSUM62. This distance is simply the sum of the number of vertices on the GO hierarchy from two annotations to their deepest common ancestor. While this method does not consider the notion of informativity and semantic similarity as rigorously as the GOGO algorithm, it gives an easily interpretable picture of how close our new annotations are to their relatives in the baseline annotation set and how often they are deeper in the GO hierarchy.

## Supporting information

Experimentally-validated functions.tsv

Non-validated Functions with High Confidence

Non-validated Functions with Low Confidence

## Acknowledgement

We gratefully acknowledge the support of NIH grant R01GM127701.

## Footnotes

1 One nat is the amount of information contained by knowing the state of a variable that has a 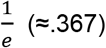 probability of occurrence.

