## Supplementary figures and images for "New Functions Identified for HIV gp120 with the New ProtSub Sequence Matching"

### Non-validated Functions with High Confidence

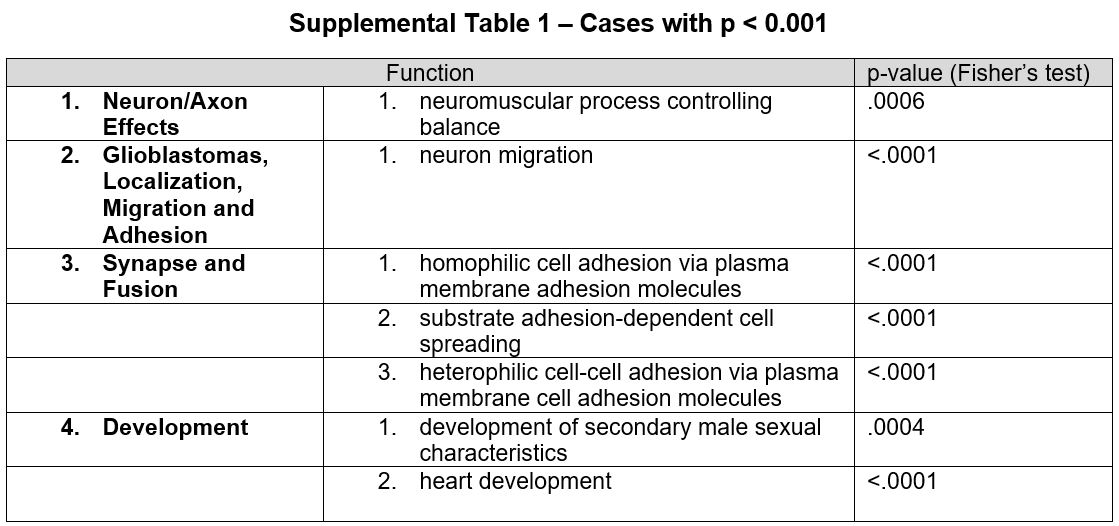

### Non-validated Functions with Low Confidence

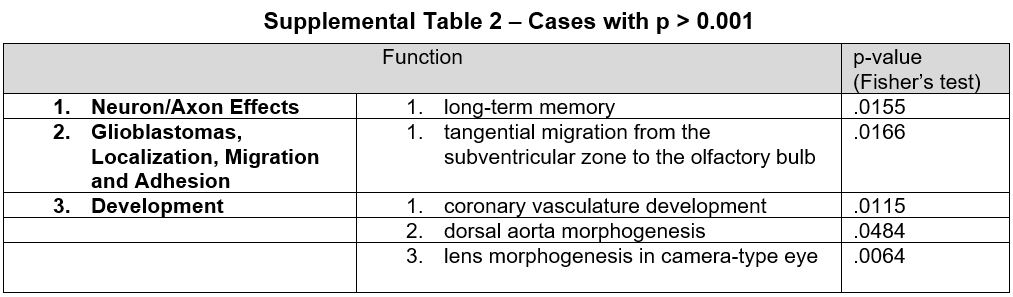
